# Signatures of heuristic-based directed exploration in two-step sequential decision task behaviour

**DOI:** 10.1101/2023.05.22.541443

**Authors:** A. M. Brands, D. Mathar, J. Peters

**Affiliations:** Biological Psychology, Department of Psychology, University of Cologne, Germany

**Keywords:** computational psychiatry, model-based, exploration, two-step task, neurocomputational endophenotypes

## Abstract

Processes formalized in classic Reinforcement Learning (RL) theory, such as model-based (MB) control and exploration strategies have proven fertile in cognitive and computational neuroscience, as well as computational psychiatry. Dysregulations in MB control and exploration and their neurocomputational underpinnings play a key role across several psychiatric disorders. Yet, computational accounts mostly study these processes in isolation. The current study extended standard hybrid models of a widely-used sequential RL-task (two-step task; TST) employed to measure MB control. We implemented and compared different computational model extensions for this task to quantify potential exploration mechanisms. In two independent data sets spanning two different variants of the task, an extension of a classical hybrid RL model with a heuristic-based exploration mechanism provided the best fit, and revealed a robust positive effect of directed exploration on choice probabilities in stage one of the task. Posterior predictive checks further showed that the extended model reproduced choice patterns present in both data sets. Results are discussed with respect to implications for computational psychiatry and the search for neurocognitive endophenotypes.

## Introduction

**“When we remember we are all mad, the mysteries disappear and life stands explained.”**-*Mark Twain* Or at least it starts to make a whole lot more sense. The notion that mental health is an integral part to all of our lives and may vary over time on a continuous scale lies at the heart of dimensional psychiatry. Such dimensional approaches address the criticism of clear-cut categorical demarcations between mental disorders. In this view, symptoms exist on a spectrum, with sub-clinical symptom variations (e.g. of depressed mood, compulsive or avoidant behaviours etc.) present in the *healthy population* (Insel et al., 2010; Robbins et al., 2012).

Transdiagnostic research specifically tackles the traditional symptom-based categorisation and thereby partitioning of mental disorders. High rates of comorbidity present another common issue raised with regard to the current conceptualisation and point to inherent flaws (i.e. commonly co-occurring diseases might be better understood as one shared rather than two distinct entities; Insel et al., 2010; Dalgeshi et al., 2020). Oftentimes, transdiagnostic and dimensional approaches go hand in hand as they both aim to improve our understanding of mental disorders, and aim to bring these approaches more in line with syndrome-based understanding known from other medical disorders (e.g., one does not have leg pain, but a broken leg; Insel 2014; Conway & Krueger, 2021).

Research into the basic computational processes that may go awry in the case of mental disorders provides important groundwork for these efforts (Adams et al., 2016). Computational psychiatry has identified several key mechanisms which likely cut across traditional diagnostic lines (Montague et al., 2012; Huys, Maia, & Frank, 2016; Insel et al., 2010; Moutoussis, Eldar, & Dolan, 2017). Such computationally derived *transdiagnostic endophenotypes* might better differentiate between mental health and disease than symptom-based conceptualisations (Robbins et al., 2012; Wise & Dolan, 2020, Yip et al., 2022; Conway & Krueger, 2021). All of these recent changes ultimately aim at furthering our understanding of mental disorders, improving prevention, diagnosis and treatment options. The hope is to someday be able to provide *precision medicine,* more akin on the common understanding and handling of “medical” disorders (Insel, 2014).

Reinforcement Learning (RL) theory (Sutton & Barto, 2018) has been of central importance in these efforts and extensively studied (Montague et al., 2012; Huys et al., 2021; Wise & Dolan, 2020). Here, two processes have emerged as promising computational endophenotypes mapping onto (sub-) clinical variation in symptoms -*model-based* control and *exploration* behaviour (Goschke 2014; Addicot et al., 2017). Regarding model-based control, behaviour is thought to depend on at least two systems: a *model-free* (MF) system, which selects actions based on past reinforcement and a *model-based* (MB) system that uses a model of the environment to select actions based on a computation of their predicted consequences (e.g. Balleine & O’Doherty, 2010; Daw et al., 2011; Daw & O’Doherty, 2014). The exploration-exploitation trade-off (Addicot et al., 2017; Sutton & Barto, 2018) refers to the process of balancing between selecting novel courses of action (exploration) and doing what has worked in the past (exploitation; Daw et al., 2006; Gershman, 2018, 2019). Here, at least two strategies have been discussed (Gershman, 2018; Wilson et al.,2014; 2021): choice randomization (random exploration), e.g. via SoftMax or epsilon-greedy choice rules (Sutton & Barto, 2018) and directed exploration, which involves the specific selection of options that maximize information gain (Wilson et al., 2021). Interestingly, despite the fact that both model-based control and directed exploration have been conceptually linked to the goal-directed system, both processes have largely been studied in isolation.

For more than a decade, the *two-step task* (TST, Daw et al., 2011), as well as various variants thereof, have been the key paradigm in the study MB and MF contributions to behaviour in human studies. This task has also been a central instrument in computational psychiatry (e.g. Voon et al., 2017; Patzelt et al., 2019; Dolan & Dayan, 2013). The TST is a sequential RL task, consisting of two stages, which each involve a choice between two options (Daw et al., 2011, Figure 1). First-stage choices lead to one of two different second stages (S2, Figure 1) with either high probability (common transition, 70%) or low probability (rare transition, 30%). Transition probabilities are reversed for the other S1 option (Figure 1). Second stage options are then associated with either drifting reward probabilities (*classic version*, see e.g. Daw et al., 2011; Gillan et al., 2016) or drifting reward magnitudes (*modified version*, see. Mathar et al., 2022). Following the S2 choice and outcome presentation (c.f. Figure 1) a new trial begins and participants are set back to S1.

**Figure 1.**
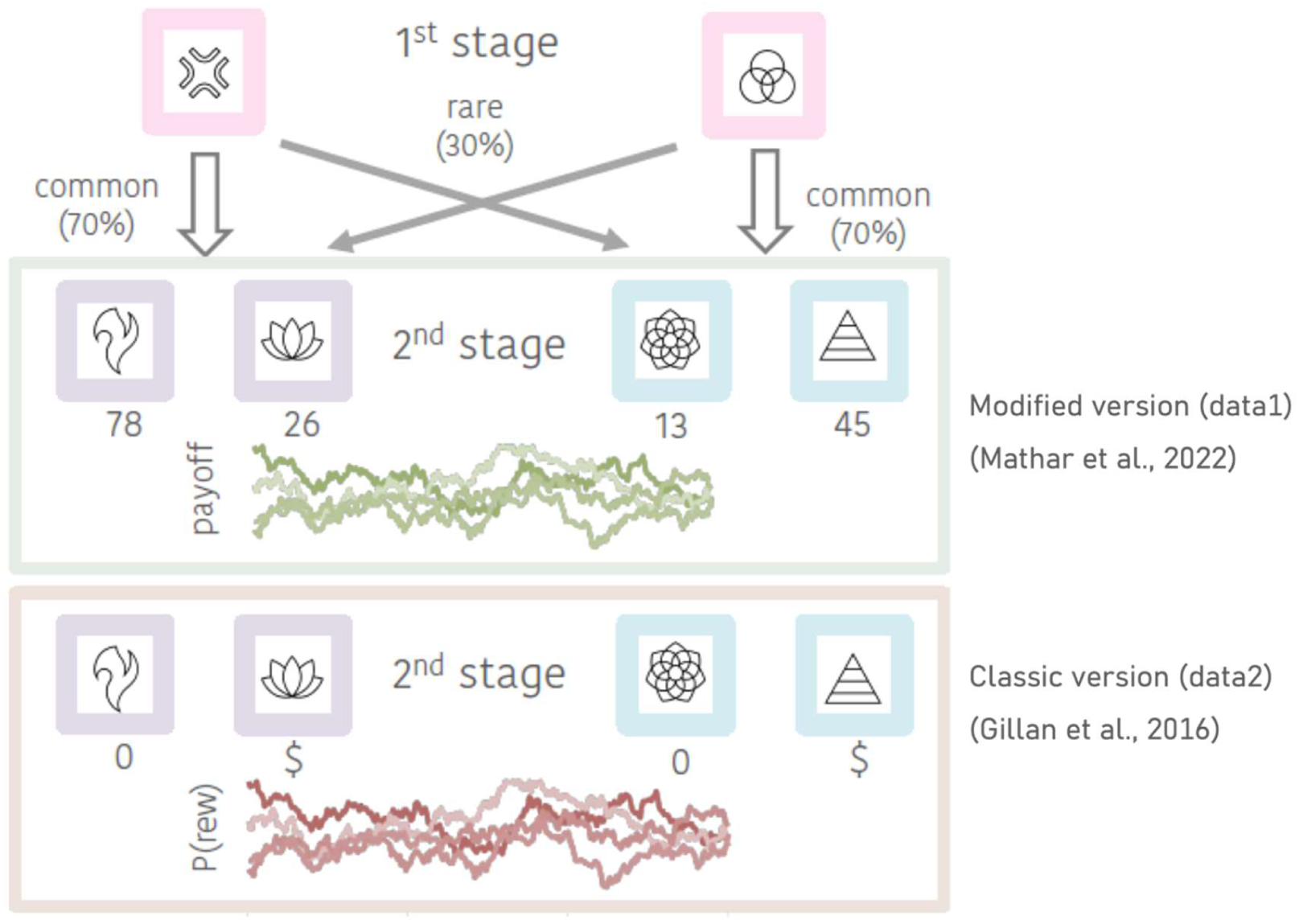
Outline of the two-step task (TST). Transition probabilities from the first stage to the second stage remain the same in both versions of the task. The second stage with a green frame depicts the modified task version employed in data set *data1* (Mathar et al., 2022): after making a S2-choice subjects receive feedback in the form of continuous reward magnitudes (rounded to the next integer). The lower S2 stage (orange frame) depicts the classic version (used in data set *data2*; Gillan et al., 2016), in which the S2 feedback is presented in a binary fashion (rewarded vs. unrewarded based on fluctuating reward probabilities).

Importantly, MB and MF control make different predictions for S1 choices as a function of reward and transition on the previous trial: a purely MF agent would repeat previously rewarded S1 actions, regardless of transition. In contrast, a MB agent “knows” about the transition structure, and would therefore switch to the other S1 option following the experience of a high reward after a rare transition, and repeat the same S1 choice following the experience of a low reward after a rare transition. In this way, the TST is thought to allow for an estimation of the relative contribution of each system (Daw et al., 2011; Otto et al., 2013a).

The distinction between MB and MF is conceptually closely related to *goal-directed* vs. *habitual,* behavioural control (Daw et al., 2006; Daw & O’Doherty, 2014; Kool, Cushman, & Gershman, 2018). Despite growing criticisms of this simplified view (see e.g. Akam et al., 2015; Collins & Cockburn, 2020; Feher da Silva & Hare, 2018), this terminology is widely used interchangeably (Miller, Shenhav, & Ludvig, 2019).

A reduction in MB control and resulting over-reliance on the MF system is thought to underlie habitual and/or compulsive behaviors characteristic of several mental disorders (e.g. Voon et al., 2015). Many studies since have indeed provided empirical support for this idea, showing reduced MB control in several patient groups spanning schizophrenia (Culberth et al., 2016), substance use disorders (e.g. Reiter et al., 2016; Sebold et al., 2017), pathological gambling (e.g. Bruder et al., 2021; Wyckmans et al., 2019), eating disorders (Foerde et al., 2021; Reiter et al., 2017), and obsessive-compulsive disorder (OCD; Gillan et al., 2011, Gillan & Robbins, 2014; Brown et al., 2020). Similar effects were observed for sub-clinical variations in symptom severity (Gillan et al., 2016; Seow et al., 2021). Reduced MB control might thus constitute a promising transdiagnostic endophenotype that might more closely relate to real world behaviour than traditional clinical categorizations (Ferrante & Gordon, 2021; Maia & Frank, 2011).

As noted above, directed exploration shares some conceptual features with MB control. Both are assumed to depend on more elaborate computations (Daw & O’Doherty, 2014; Gershman & Daw, 2012; Wilson et al., 2021; for diverging views regarding the dichotomy of MB & MF control see e.g. Akam et al., 2015; Doody et al., 2022; Miller et al., 2019). However, directed exploration might also rely on simpler heuristic-based exploration strategies (Fox et al., 2020). For example, an agent may utilize a simple proxy measure of environmental uncertainty (and therefore of potential information gain) rather than a precise model of environmental dynamics, which is often assumed in e.g. Kalman Filter models (e.g. Daw et al., 2006; Speekenbrink, 2022; Chakroun et al., 2020). Reductions in directed exploration have been observed in a range of mental disorders, including substance use disorders (SUD; Addicot et al, 2013; Morris et al., 2016; Smith et al., 2020), gambling disorder (Wiehler, Chakroun, & Peters., 2021), OCD, Depression (Blanco et al., 2013) as well as anxiety (Aberg et al., 2021; Smith et al., 2020). Similar to model-based control, aberrant directed exploration might constitute a potential transdiagnostic endophenotype in computational psychiatry (Addicot et al., 2017).

Interestingly, the TST shares several key features with tasks traditionally used in the study of directed exploration. For example, similar to restless bandit tasks used to study exploration (Daw et al., 2006; Sutton & Barto, 2018; Speekenbrink, 2022; Chakroun et al., 2020; Wiehler et al., 2021), rewards (or reward probabilities, depending on task version) in S2 of the TST drift according to Gaussian random walks. Potential information gain in S2 of the TST is therefore a function of sampling recency of S2 options. Given the structural similarities between S2 of the TST and restless bandit problems, it thus seems natural to hypothesize that similar directed exploration processes might contribute to TST behaviour. However, there have only been isolated attempts to integrate these concepts (Gijsen et al., 2022). A further issue relates to recent concerns regarding the interpretation of TST results. Feher da Silva and Hare (2018; 2020) argued that MB measures from the TST only reflect one specific model-based strategy (see also Toyama et al., 2017;2019). Participants might well utilize different and/or additional models of the task that constitute a “model-based” strategy, but which are not directly measured using the simple MB vs. MF dichotomy embedded in standard hybrid models (Collins et al., 2017; Collins & Cockburn, 2020; Feher da Silva et al., 2022). To resolve this issue, some researchers have resorted to specifying task features and experimental procedures which allow for a more precise interpretation of results from classical analyses (see e.g. Kool et al., 2016). Another way of tackling this issue is to specify and implement proposed additional processes within computational models, which can subsequently be tested with regard to their fit to empirical data.

The present study follows the latter approach. We had the primary aim to extend standard hybrid models of MB and MF control in the TST by incorporating directed exploration mechanisms. To this end, we implemented and compared several potential candidate mechanisms, including more elaborate mechanisms of uncertainty tracking (Kalman, 1960; Daw et al., 2006) as well as simpler heuristics (Fox et al., 2020). We tested and compared our models in two independent data sets, a variant of the TST with drifting reward magnitudes in S2 (*data1*, Mathar et al., 2022) as well as the classical TST with drifting reward probabilities (and binary payouts) in S2 (*data2*, Gillan et al., 2016). A wealth of empirical data from the TST have already been acquired, many in clinical groups. These data are often available for re-analysis. Therefore, investigations into additional computational mechanisms that might be reflected in these data could proof valuable for the field.

## Methods

### Participants and Task Versions

We evaluated all models on the basis of a re-analysis of two independent existing data sets. The first data set (*data1*) encompasses data from 39 healthy, male participants (aged 18-35; *M*= 25.17, *SD*= 3.89) who performed 300 trials of the modified TST version (neutral condition from Mathar et al., 2022). The second data set (*data2*) constitutes a subsample (N=100) from a previously published large scale online study using the classical TST (Gillan et al., 2016).

### Data1

The first data set (data1) was obtained in a recent study (Mathar et al., 2022) that spanned two testing sessions and included additional tasks, self-report measures and physiological markers of autonomic arousal, which were not analysed for the current study (for more details see Mathar et al., 2022). Prior to performing the TST, participants received instructions regarding the transition probabilities as well as fluctuating reward structure and performed 20 training trials. In addition, they were informed that the maximum response time was two seconds and that they could obtain an additional 4€ in reimbursement, contingent upon task performance.

This TST version used transition probabilities fixed to 70% and 30% for common and rare transitions, respectively and the reward magnitudes for each S2 option followed independent Gaussian Random Walks (fluctuating between 0 and 100, rounded to the next integer, see Fig. 1). S2 states that were marked by different colours to make them more easily distinguishable. For all analyses, trials with response times < 150ms were excluded.

### Data2

Gillan and colleagues (2016) used the original variant of the TST (Daw et al., 2011). Here, reward probabilities of all choice options varied independently according to Gaussian Random Walks, and participants received binary reward feedback (rewarded vs. no reward; Fig.1). Detailed descriptions of the whole sample, exclusion criteria, procedure, additional measures, as well as specifics of the TST version employed can be found in the original publication.

We drew a subsample of N=100 (age: M=34; SD=11; 69% female) from the full sample of Experiment 1 from the original publication (N=548, age: M=35; SD=11; 65% female; for further details see Gillan et al., 2016). This subsample is representative of the whole original sample regarding the self-report measures obtained by the authors. To yield this subsample, we randomly sampled from the original full sample from Experiment 1 until self-reported symptom severity did not significantly differ from those of the full sample. The resulting transdiagnostic symptom-score relating to compulsivity and intrusive thoughts (c.f. PCA analyses reported in Gillan et al., 2016 for more detail) was chosen as a criterion due to its significant association with model-based RL in the original publication.

### Model-Agnostic Analyses

As a first step we used logistic mixed effects regression models to analyze *stay-probabilities* for first-stage choices, i.e. the probability to repeat the first stage selection of the preceding trial, depending on the transition (common vs. rare) and reward (rewarded vs. unrewarded) experienced. Such regression models are amongst the most common ways of analyzing TST behavior outside a computational modeling framework. Reward and transition type (rewarded/unrewarded and common/rare were coded as 1/-1 respectively) were entered as (fixed effects) predictors of S1 choice repetition (i.e. perseveration). Individual subjects were entered as random effects. Using the *lme4* package (Bates et al., 2015) this resulted in the following model specification in the R syntax:

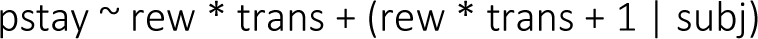

Due to the presentation of continuous reward magnitudes in the modified version (data1) we defined outcomes to be *rewarded / unrewarded* (1/-1) relative to the mean outcome over the preceding 20 trials (Wagner et al., 2020; Mathar et al., 2022).

As additional model-agnostic indices of MB and MF behavior, we calculated difference scores (*MBdiff & MFdiff)* as proposed by Eppinger and colleagues (2013):

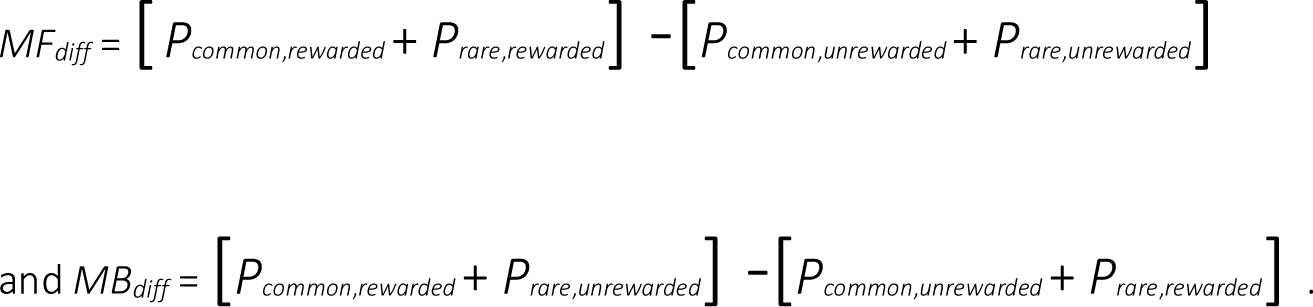

### Computational Models

Our model space consisted of nine models in total. Two standard hybrid RL models without exploration terms but with different learning rules served as baseline models (Q-Learner, Bayesian Learner). We extended both models with exploration terms that incorporated different ways in which S2 uncertainty might impact first stage choice probabilities (see below).

### Learning Rules

#### Q-Learner

The first model is an adaptation of standard hybrid models (e.g. Daw et al.,2011; Otto et al., 2013b). Here, for each first-stage state-action pair (i.e. choice option), separate MF and MB values (𝑄_𝑀𝐹_, 𝑄_𝑀𝐵_) are calculated in parallel.

MF values in both stages are updated using the TD learning algorithm SARSA (Rummery & Niranjan, 1994), such that MF Q-values of a chosen state-action pair at stage 𝑖 in trial 𝑡 are updated according to:

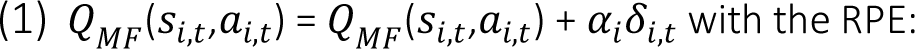

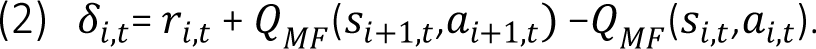

and a constant learning rate 𝛼_i_(ranging from 0 to 1) for each stage.

S2 prediction errors are incorporated into S1 value estimates via the second-stage learning rate 𝛼_2_ (constrained between 0 and 1):

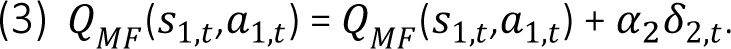

While several other models utilize an *eligibility trace* parameter (𝜆) to propagate S2 RPEs, here we chose to reduce the parameter space and model complexity by instead using the S2 learning rate. This formalization was used for all QL models, while all BL models kept the an additional parameter 𝜆 (as there is no 𝛼_2_, for more detail see below; c.f. Table 1).

**Table 1.** Free and fixed parameters for all models.

| Model | free parameters | fixed parameters |
| --- | --- | --- |
| M1 (BL) | $\alpha, \alpha_3, \lambda, \beta_{MB}, \beta_{MF}, \beta_C, \beta_2$ | $\hat{\gamma}, \hat{\theta}, \hat{\sigma}_o, \hat{\sigma}_d, \hat{\sigma}_{n,1}^{2pre}, \hat{\mu}_{n,1}^{pre}$ |
| M2, M4, M6 (BL + EB) | $\alpha, \alpha_3, \lambda, \beta_{MB}, \beta_{MF}, \beta_C, \beta_2, \phi$ | $\hat{\gamma}, \hat{\theta}, \hat{\sigma}_o, \hat{\sigma}_d, \hat{\sigma}_{n,1}^{2pre}, \hat{\mu}_{n,1}^{pre}$ |
| M8 (BL + EB<sub>bandit</sub> + EB<sub>trial</sub>) | $\alpha, \alpha_3, \lambda, \beta_{MB}, \beta_{MF}, \beta_C, \beta_2, \phi, \rho$ | $\hat{\gamma}, \hat{\theta}, \hat{\sigma}_o, \hat{\sigma}_d, \hat{\sigma}_{n,1}^{2pre}, \hat{\mu}_{n,1}^{pre}$ |
| M3 (QL) | $\alpha, \alpha_2, \alpha_3, \beta_{MB}, \beta_{MF}, \beta_C, \beta_2$ | |
| M5, M7 (QL + EB) | $\alpha, \alpha_2, \alpha_3, \beta_{MB}, \beta_{MF}, \beta_C, \beta_2, \phi$ | |
| M9 (QL + EB<sub>bandit</sub> + EB<sub>trial</sub>) | $\alpha, \alpha_2, \alpha_3, \beta_{MB}, \beta_{MF}, \beta_C, \beta_2, \phi, \rho$ | |

*Note.* QL refers to the basic hybrid model. BL denotes the altered hybrid model versions with Kalman Filter updates. E = added first-stage exploration bonus; $\phi$ : parameter that scales the exploration bonus. Note that this parameter remains the same for both exploration bonus variants, regardless of the specific formalisation of uncertainty estimates in a given model. For M8 and M9 an additional free parameter $\rho$ was included to account for additive exploration effects of both heuristics.

We included an additional “forgetting process” for MF Q-values (Toyama et al., 2017, Toyama et al., 2019), such that unchosen Q-values decayed towards the mean according to a decay rate 𝛼_3_(constrained between 0 and 1):

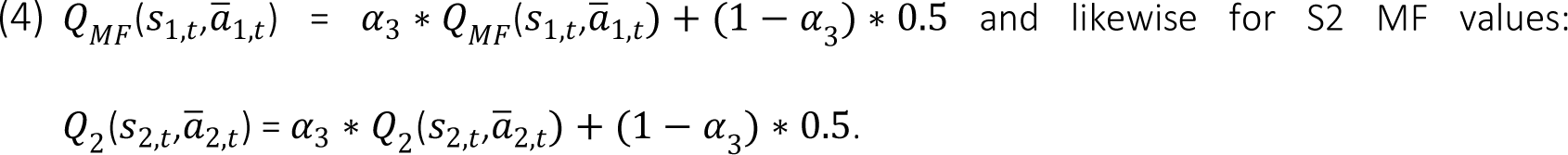

Recall that transition probabilities in the model were fixed as follows:

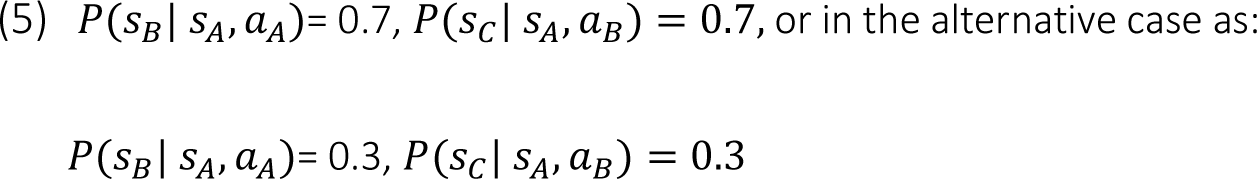

with

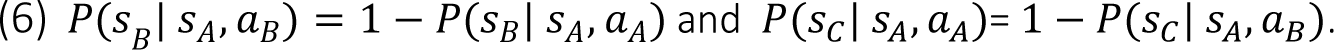

First-stage 𝑄_𝑀𝐵_values were then computed as the maximal 𝑄_2_values weighted by their respective transition probabilities. Thus, using the Bellman equation 𝑄_𝑀𝐵_ values are defined as:

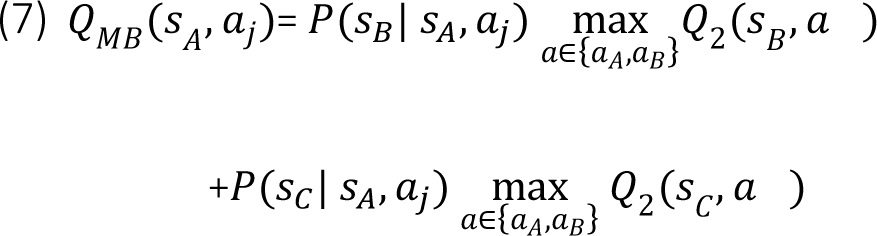

As a trial ends with the second-stage choice, for S2 only MF values are relevant, such that:

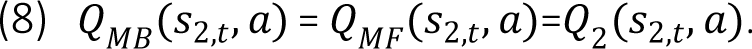

Accordingly, 𝑄_2_(𝑠_2,𝑡_, 𝑎) updates follow the TD process as described previously for first stage 𝑄_𝑀𝐹_ values (Equation 1), while allowing for a separate learning rate 𝛼_2_ (also constrained between 0 and 1).

#### Bayesian Learner (BL)

The second learning rule applied is based on the *Bayesian Learner* (BL) as commonly applied in restless bandit tasks (Daw et al., 2006; see also: Chakroun et al., 2020; Wiehler et al., 2021). Here, the constant learning rate 𝛼_2_ (see Equation 1) is replaced by a trial-specific learning rate (𝜅_𝑡_) based on the Kalman Filter (Kalman, 1960). The idea is that participants track the changes in the underlying reward means of all choice options, as well as the uncertainty associated with these estimates. In this model, learning rates are then uncertainty-dependent.

For both data1 and data2, the true process underlying the Gaussian random walks of S2 rewards was directly incorporated in the model (for applications in the explore-exploit dilemma see e.g. Daw et al., 2006; Chakroun et al., 2020).

For data1, rewards for each option 𝑛 at trial 𝑡 ranged between 0 and 1 (multiplied by 100 and rounded to the next integer for presentation as points in the task; c.f. Methods). These were generated following a Gaussian random walk with mean 𝜇_n,t_ = 0.5, and SD = 0.04 (observation variance 𝜎^2^_*o*_ = 0.04^2^). Means for each option independently changed on a trial-to-trial basis.

As participants were assumed to represent the reward-walk dynamics in their internal models, the computational model includes fixed parameters γ̂, 𝜃, 𝜎̂_𝑜_, 𝜎̂_𝑑_, which are set to values approximating those of the underlying random walk: the decay parameter 𝛾 = 0.9836, decay centre 𝜃= 0.45, and the diffusion variance 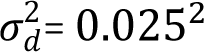, and diffusion noise 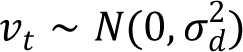.

Participants start with prior beliefs of a normally distributed reward mean 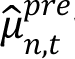 with variance 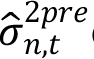 of a chosen second-stage option 𝑛 on trial 𝑡 and update these in light of the reward 𝑟_t_ they receive according to:

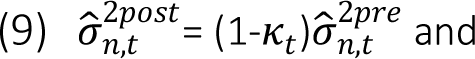

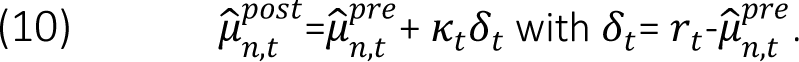

The parameter 𝜅_𝑡_ is the Kalman Gain, which serves as the uncertainty-dependent learning rate, similar to 𝛼_2_ in the original (QL) model. In the updating process of a chosen option the Kalman Gain (just like 𝛼) scales the prediction error used for updating the mean reward estimate 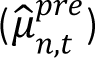. The important difference being that 𝜅_𝑡_ varies on a trial-to-trial basis depending on the observation as well as diffusion variance:

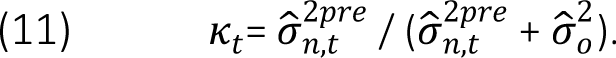

Thus, high observation uncertainty yields large values of 𝜅_t_ and thus increased updating. In contrast, low observation uncertainty yields small values of 𝜅_𝑡_ and thus reduced updating.

Values of all options are updated between trials according to:

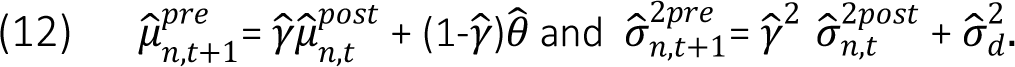

That is, between-trial updating describes participants tracking of the assumed dynamics underlying the Gaussian random walks. Due to these between-trial updating dynamics, the forgetting process for S2 Q-values (see Equation 4) was omitted from all BL models.

The Kalman Filter (Equation 12) was implemented as the learning rule for second-stage values and then incorporated into the hybrid model, such that 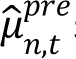 substitutes the Q-values for second-stage options in Equation 13 above.

### Choice Rules

The standard SoftMax function (*SM,* McFadden, 1973; Sutton & Barto, 2018) served as the basis for all choice rules. According to this rule, choices’ probabilities scale with the value differences between options.

#### SoftMax

As proposed by Otto and colleagues (2013b), separate coefficients for 𝑄_𝑀𝐹_ and 𝑄_𝑀𝐵_ were used, rather than a single weighting parameter 𝜔 (as done e.g. in Daw et al., 2011 and Gillan et al., 2016). Thus, choice probabilities for action 𝑎 at the first stage were modelled as:

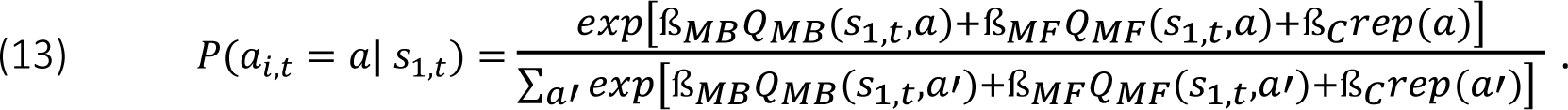

The parameter ß_𝐶_ describes the “stickiness” of first stage choices, i.e. first order perseveration. The indicator function 𝑟𝑒𝑝(𝑎) equals 1 if the first-stage choice of the previous trial is repeated and 0 otherwise.

At the second stage, choices are driven by MF Q-values only (𝑄_2_(𝑠_2,𝑡_, 𝑎), as described above) scaled by the second-stage inverse temperature parameter ß_2_, such that:

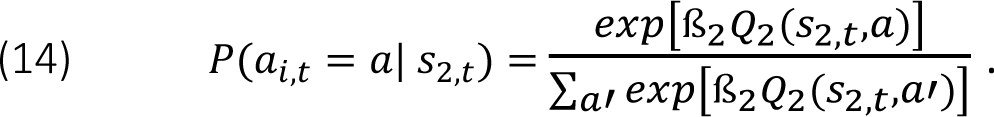

The two baseline models M1 and M3 (BL & QL respectively) used this basic version of the SM. These were extended by incorporating terms that account for directed exploration in first stage choices.

#### Exploration Bonus (eb)

The following sections describe the different implementations of directed exploration for first stage choices in more detail. Generally, we compared various implementations of an exploration bonus (𝑒𝑏) for first-stage values to capture strategic exploration. The general idea is that participants may seek out uncertain S2 states for information-gain and potential long-term reward maximization. Random exploration, in contrast, is assumed to result from sub-optimal, random deviations from a reward-maximizing decision-scheme (Sutton & Barto, 2018; Wilson et al., 2021).

For all variants of the exploration bonus 𝑒𝑏, this bonus was included in the standard SoftMax and weighted by an additional free parameter 𝜙, resulting in the following first-stage choice probabilities (for models M2 and M4 to M7):

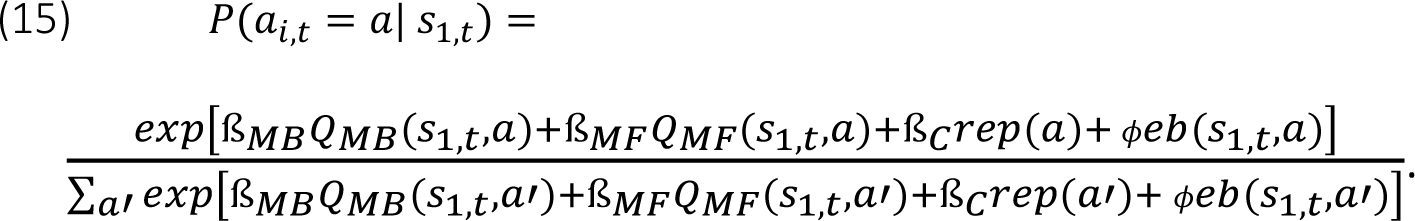

Models M8 and M9 included two measures of *eb*, therefore another free parameter 𝜌 was included modelling 𝑒𝑏_trial_ in addition to 𝑒𝑏_bandit_, resulting in:

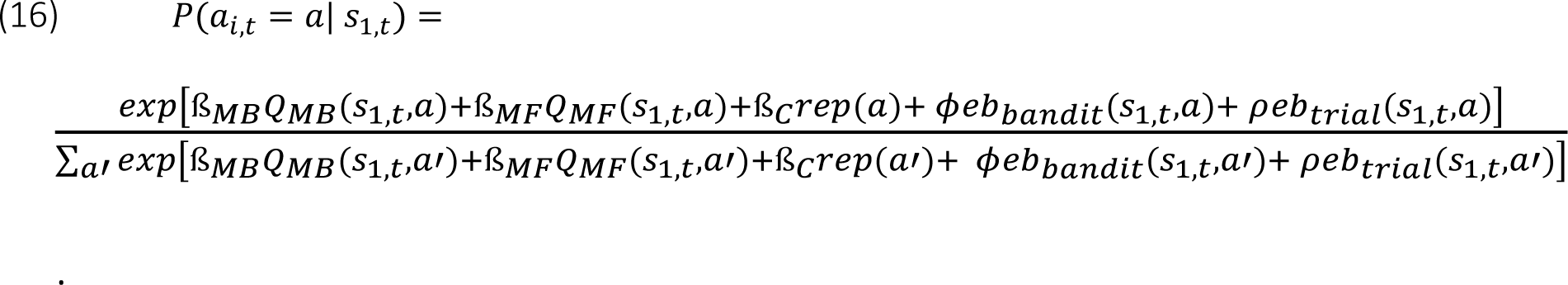

Overall, we compared four different formalizations of uncertainty that participants might draw upon during directed exploration for S1 decisions (see below). These different types of exploration bonus incorporated transition probabilities analogously to the 𝑄_𝑀𝐵_ values (c.f. Eq. 7 & Eq. 18-20). This formalization was based on previous research efforts providing evidence for the demarcation of at least two separate exploration strategies (Sutton & Barto, 2018; Wilson et al., 2014). Directed (vs. random) exploration is assumed to reflect an effortful, goal-directed strategy, aiming at long-term reward accumulation via maximal information gain. In this way directed exploration and MB control show a large conceptual overlap (i.e. deliberate forward-planning under consideration of environmental dynamics, with a long-term perspective on goal-attainment). Consequently, formalizations of directed exploration in model variants presented here were defined analogous to the MB component. In both components transition probabilities are utilized to weigh the reward and uncertainty estimates (MB and exploration, respectively) associated with S2 options to reflect these assumed deliberate and foresighted aspects.

#### Uncertainty estimates based on the Kalman filter

This variant was only included for the BL model (M2), as the Q-learning models lack an explicit representation of uncertainty. Here, the 𝑒𝑏 was based on the estimated maximum standard deviation of second-stage options (Daw et al., 2006; Chakroun et al., 2020) such that:

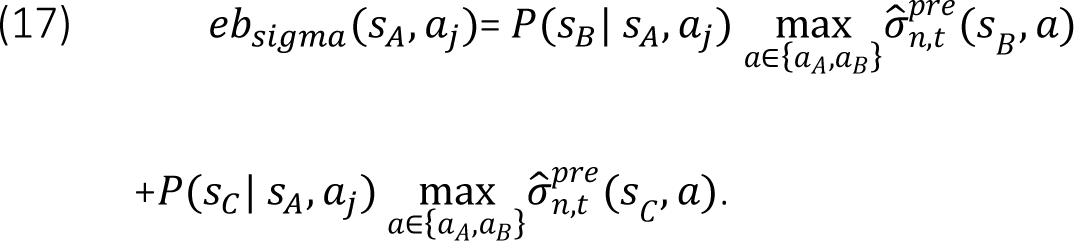

The resulting first-stage choice probabilities were modelled according to Equation 15.

All remaining variants of the exploration bonus were based on simpler counter-based heuristics.

#### Uncertainty estimates based on a bandit-counter heuristic

For the first of these (𝑏_𝑛,𝑡_(𝑠_𝐵_, 𝑎), *bandit-heuristic)*, participants were assumed to estimate how many of the alternative S2 options they have sampled since last choosing a given option 𝑛. Following selection of an S2 option 𝑛, the respective counter 𝑏_𝑛,𝑡_ is reset to 0. Thus, 𝑏_𝑛,𝑡_ of a given S2 option 𝑛 ranges from 0 (this option was chosen on the last trial) to 3 (all other S2 option were sampled since last sampling this option). Diverging from the formalization in the *BL + sigma* model (M2), here we assumed participants to sum up these counters over both options of the respective S2 stage associated most likely with either first-stage choice (instead of tracking the maximum). The sum of 𝑏_𝑛,𝑡_ across associated S2 options were again weighted by their transition probabilities (see M2; Equation 17) resulting in:

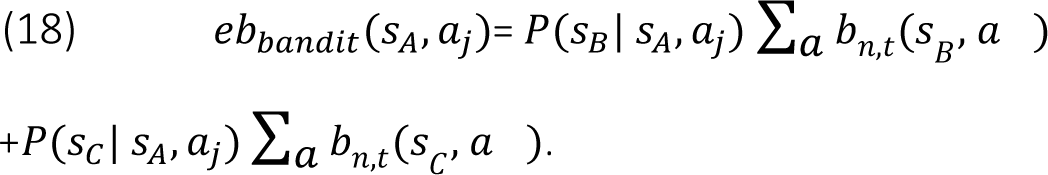

This yielded the models M4 (*BL+ bandit*) and M5 (*QL + bandit;* see Equation 15 for corresponding S1 choice probabilities). Often times uncertainty sum-scores are associated with less goal-directed exploration strategies (i.e. random exploration based on total uncertainty), such as Thompson Sampling. In such cases however, total uncertainty scores (sums) are directly linked to choice stochasticity (c.f. Gershman, 2018, 2019; Fox et al., 2020). In contrast, here the summed uncertainty proxy is incorporated in a more complex model of a decision sequence (exploration boni are sensitive to the transition type and are ultimately assigned to S1 state-action pairs). In this way higher sum scores of given counters associated with a particular S1 action are incorporated in a simplified, yet still model-based way. Moran and colleagues (2019) have furthermore used similar formalizations to describe interactive dynamics and partial overlap of the proposed MB and MF system. The authors provide evidence for the incorporation of rather parsimonious MF-like value estimates (via sum scores) to retrospectively assign credit to previous actions using an internal model of the environment.

#### Uncertainty estimates based on a trial-counter heuristic

For models *BL+ Trial* and *QL + Trial* (M6 and M7, respectively) we followed the same logic, with the only difference being that participants were assumed to utilize a trial counter (𝑡_n,t_) as a proxy for uncertainty. This counter heuristic was simply defined as the number of trials since that particular second-stage option was last sampled. The resulting exploration bonus was thus defined as follows:

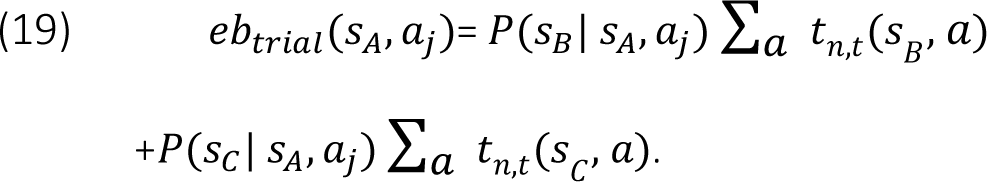

Analogous to models M4 and M5, action probabilities for first-stage choices were modelled according to Equation 15.

#### Combined models

Finally, combined models with terms for both trial- and bandit-heuristic exploration were examined, i.e. *BL+ Trial + Bandit* and *QL + Trial + Bandit* (M8 and M9, respectively). The implementation of both exploration mechanisms resulted in first-stage action probabilities according to Equation 16.

### Hierarchical Bayesian Modelling Scheme

Table 1 provides an overview of all free and fixed parameters for the models described above. Using a hierarchical Bayesian modelling scheme, subject parameters were drawn from shared group-level Gaussian distributions. This resulted in two additional free parameters 𝑀^𝑥^and 𝛬^𝑥^ for each subject-level parameter 𝑥. Group-level parameter means (𝑀^𝑥^) were assumed to be normally distributed (*M*= 0, *SD=* 10) and standard deviations (𝛬^𝑥^) were set to follow a uniform distribution (with limits 0 and 10 for 𝜆 and all 𝛼_𝑖_, and an upper limit of 20 for remaining group-level SD parameters). All learning rates (𝛼_1_, 𝛼_2_, 𝛼_3_) as well as the eligibility trace parameter (𝜆) were then back-transformed to the interval (0,1) using STANs built in cumulative density function. This was done directly within the model, so that raw subject-level parameter values ranging from -10 to 10 were mapped onto the interval (0,1).

For models M1, M2, M4, M6 and M8, participant’s estimates of the random walk parameters (γ̂, 𝜃-, 𝜎̂_𝑜_, 𝜎̂_𝑑_) were fixed to values reasonably approximating the underlying dynamics. The estimated mean and standard deviation for the reward distribution were initialised at the first trial with 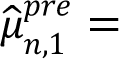 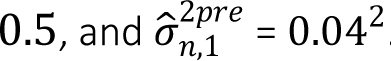.

All models were implemented using the *STAN* modelling language (version 2.21.0; Stan Development Team, 2019) running in the statistical program R, which was also used for all further analyses (version 3.6.1, R Core Team, 2019). The sampling for each model was done using a Markov-Chain-Monte-Carlo (MCMC) algorithm (no-U-turn sampler NUTS), with four chains running 10000 iterations each, 8000 of which were discarded as warm-up. MCMC methods are based on the generation of a random number sequence (chain) that is used to sample a probability distribution. Parameters estimates with higher (posterior) probability are sampled more often, resulting in a posterior probability distribution. The desired state is reached when a chain has reached equilibrium. 𝑅^-^ is a measure of convergence across chains, indicating the ratio of between-chain to within-chain variance. Here, values of 𝑅^-^ ≤1.1 were considered acceptable.

In a first step, all models were compared regarding their predictive accuracy. As this method only provides a relative comparison between models, posterior predictive checks were performed to gain a deeper understanding. These allow a more detailed insight with regard to predictions made by the model and their ability to accurately portrait the data as well as an indication of possible model misspecifications (Wilson & Collins, 2019).

#### Open Code

STAN model code will be shared via the *Open Science Framework* upon publication. By making the model code freely available, we wish to facilitate further application and development of this model and further adaptations thereof. Transparently reporting on model specifications also holds the potential of direct comparisons of parameter estimates (e.g. as reported above for the data sets compared here).

#### Open Data

Both data sets analyzed here are freely available (for links see the respective publications).

##### Model Comparison

Model comparison was performed using the *loo* package (Vehtari et al., 2023) which provides a measure of predictive accuracy via leave-one-out cross-validation (LOO; Vehtari, Gabry, & Gelman, 2017). To this end the estimated log pointwise predictive density (-elpd) is applied as the criterion of interest. The model with the lowest -elpd score was selected as the best fitting model. While lower values indicate a superior fit, in direct comparisons between models values close to 0 indicate superior fit. As can be seen in Table 3, in these cases the difference in elpd compared to the winning model (*elpd diff*) is provided (thus, values close to zero show “smallest” distance to the winning model). In cases in which a more parsimonious model showed overlap with the best-fitting model in terms of the SE of the elpd difference, the more parsimonious model was chosen.

## RESULTS

### Model-Agnostic analyses

In a first step, we quantified MF and MB contributions using common model-agnostic procedures (see e.g. Daw et al., 2011; Otto et al., 2013a; Gillan et al., 2016) as outlined in the methods section. A linear mixed model of S1 stay-switch behavior using the factors reward and transition type as well as their interaction confirmed the standard effects (Daw et al., 2011; Otto et al., 2013a): in both data sets (see Table 2) we observed a main effect of reward (reflecting MF control) and a reward x transition interaction (reflecting MB control).

**Table 2.** Results from regression analyses of S1 choice repetition probability.

|  |  | Estimate | 95% CI | z-Value | p-Value |
| --- | --- | --- | --- | --- | --- |
| data1 | Intercept | 1.35 | [1.12; 1.58] | 11.48 | <.01 |
|  | Reward | 0.11 | [0.05; 0.18] | 3.47 | <.01 |
|  | Transition | -0.07 | [-0.14; -0.01] | -2.14 | .03 |
|  | Reward*Transition | 0.47 | [0.36; 0.59] | 8.10 | <.01 |
| data2 | Intercept | 1.73 | [1.53; 1.94] | 16.68 | <.01 |
|  | Reward | 0.64 | [0.51; 0.77] | 9.89 | <.01 |
|  | Transition | 0.02 | [-0.03; 0.08] | 0.81 | .42 |
|  | Reward*Transition | 0.16 | [0.07; 0.24] | 3.76 | <.01 |

Note. Reward: main effect of reward type (unrewarded vs. rewarded), commonly interpreted as an indicator for MF control; Transition: main effect of transition type (rare vs. common); Reward\*Transition: interaction of Reward and Transition type, commonly interpreted as an indicator for MB control.

However, regression results along with visual inspection (Table 2, Figure 2) also suggest differences between task versions (i.e. between data1 and data2). The MF effect was somewhat more pronounced in the data set from Gillan and colleagues (2016; data2), while data1 showed a more pronounced MB effect. This contrast between data1 and data2 was also clearly evident in the respective model-agnostic difference scores for the two effects (Figure 2, lower panel).

**Figure 2.**
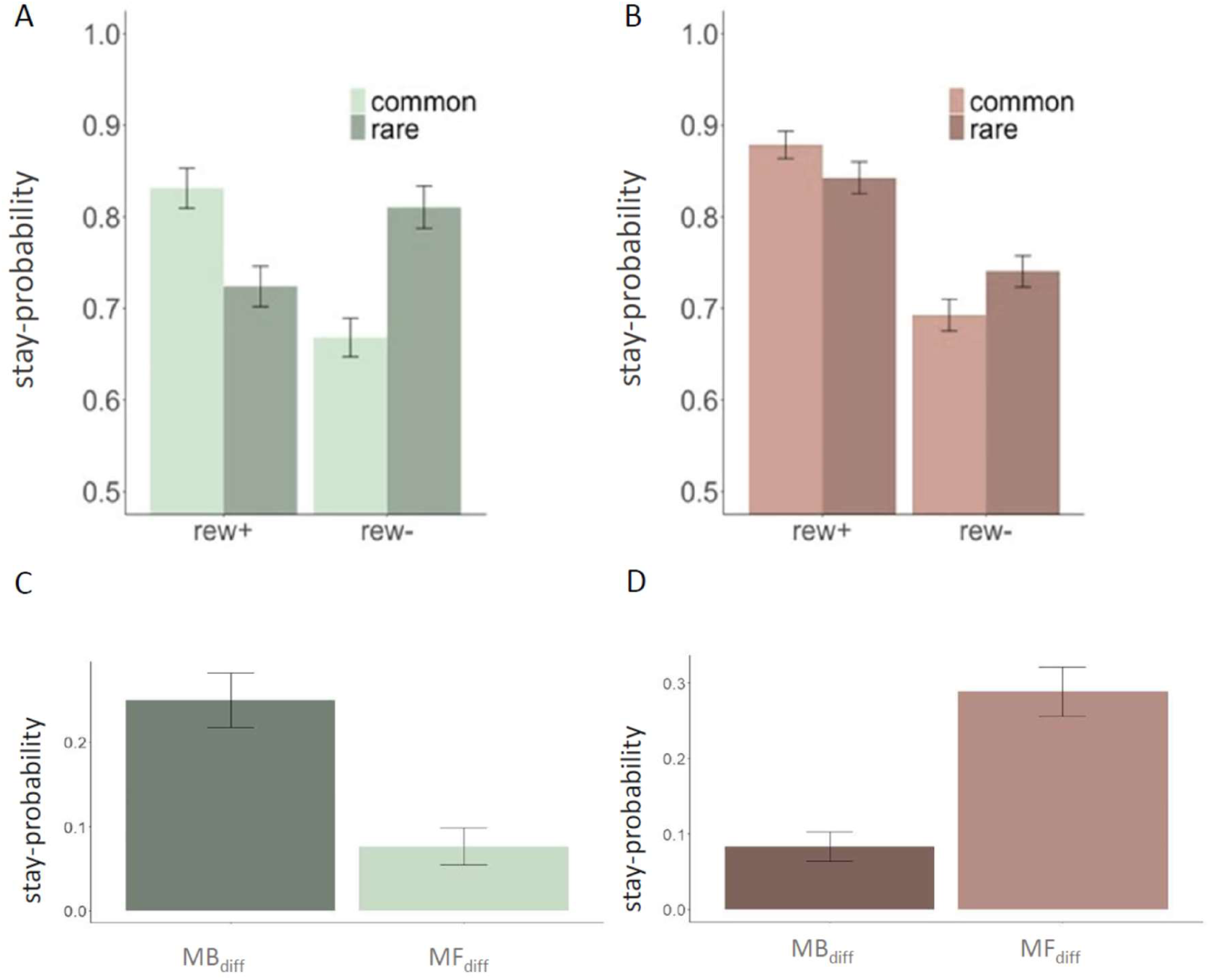
Stay-Probabilities of S1 choices and difference scores. Upper panel: Probabilities for S1 choice repetition as a function of reward (rew+ : rewarded; rew- : unrewarded) and transition type (common/rare) of the preceding trial. Lower panel: MB and MF difference scores as defined by Eppinger et al. (2013), bar heights depict mean scores over all participants, error bars show the standard error. The left plots (green, A & C) shows results from data1; the right plot (orange) shows results from data2.

### Model comparison

In both data sets all parameters (group-as well as subject-level) could be estimated well, as evidenced by the aforementioned convergence measure 𝑅^-^ (all 𝑅^-^ <1.1). Model comparison then was based on the estimated log pointwise predictive density (-elpd). Here, lower absolute values reflect a superior fit. The difference in -elpd (*-elpd diff*; c.f. Table 3) is provided in reference to the winning model (which itself thus always has a *-elpd diff* of zero). The Q-Learner models consistently outperformed BL models across both data sets (cf. Figure S1).

**Table 3.** Results from model comparison of QL-models using Leave-one-out cross-validation (LOO).

| Data Set | Model | *-elpd diff* | *se diff* |
| --- | --- | --- | --- |
| data1 | M3 (Base) | -7.9 | 6.3 |
|  | M5 (Base + bandit) | -0.4 | 5.0 |
|  | M7 (Base + trial) | -5.4 | 4.5 |
|  | M9 (Base + bandit + trial) | 0.0 | 0.0 |
| data2 | M3 (Base) | -10.8 | 8.1 |
|  | M5 (Base + bandit) | 0.0 | 0.0 |
|  | M7 (Base + trial) | -8.5 | 7.3 |
|  | M9 (Base + bandit + trial) | -4.2 | 6.9 |

*Note.* The difference in the expected log pointwise predictive density (*elpd diff*) and standard error of the difference (*se diff*). These values show the results of a model comparison using LOO estimates. Each model is compared to the preferred model (M5/M9), as there is no difference between the winning model and itself, values in the first column are always zero.

Within the group of QL models the bandit model (M5) provided the best fit for data2 and was indistinguishable from the combined model (M9) for data1 (i.e., the standard error of the elpd-difference of M5 from the best model M9 included zero, see Table 3, Figure 3). Thus, all further analyses focused on the more parsimonious model (M5, *QL+Bandit*).

**Figure 3.**
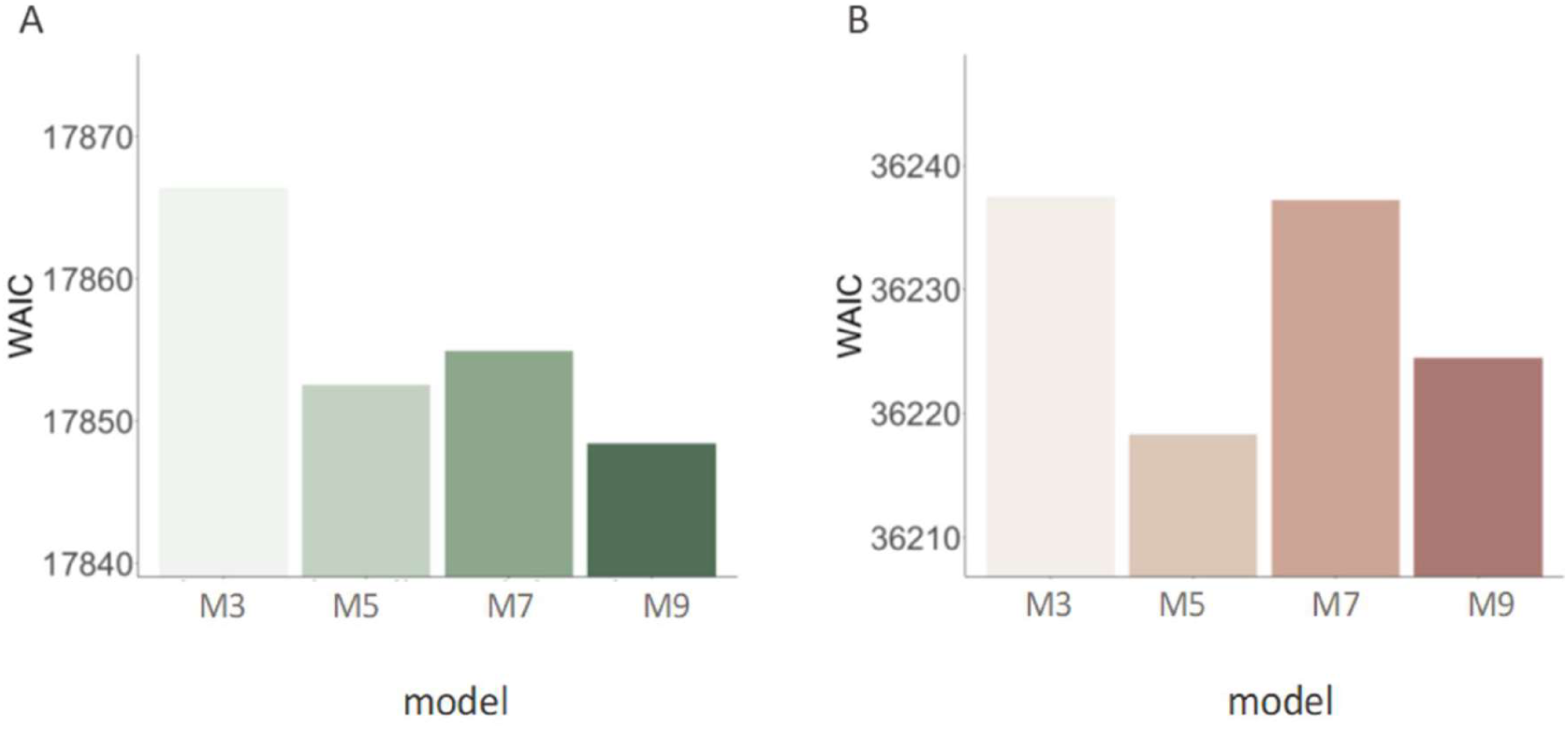
Model Comparison Results via the Widely Applied Information Criterion (WAIC) for all QL Models (M3, M5, M7, M9). Panel A/B green/orange bar plots refer to data1 and data2, respectively. M3: *QL base* model without added exploration bonus. M5/7: *Base + Bandit/ Trial* refer to the model variants with added heuristic-based exploration bonus using stimulus identity/recency, respectively. M9: *Base + Bandit+ Trial*: model variant with additive combination of both heuristic-based exploration implementations.

### Posterior Predictive Checks

While results from model comparisons can provide relative support for one computational account over another, posterior predictive checks are required to ensure that a model can reproduce core patterns in the data. We thus simulated 10000 full trial sequences, 8000 of which were discarded as warm-up, yielding 8000 (2000 per chain) simulated data sets per subject. Simulations were performed during fitting the model to the empirical data (i.e. on the basis of possible parameter values sampled during this procedure). As can be taken from Table 4, simulated S1 choices largely reproduced the patterns observed in human data. We repeated the model-agnostic analyses of stay-/switch behavior (shown above for empirical data, c.f. Table 2 Figure 2 and 4) for the simulated data sets. Visual inspection (Figure 4) revealed an underestimation of *“stay”*-probabilities in the simulations when compared to the empirical data. Nonetheless, the overall pattern of stay-switch tendencies as a function of reward and transition were largely reproduced, whereas this was less pronounced for the main effect of reward (MF contribution; Table 2, Figure 2).

**Figure 4.**
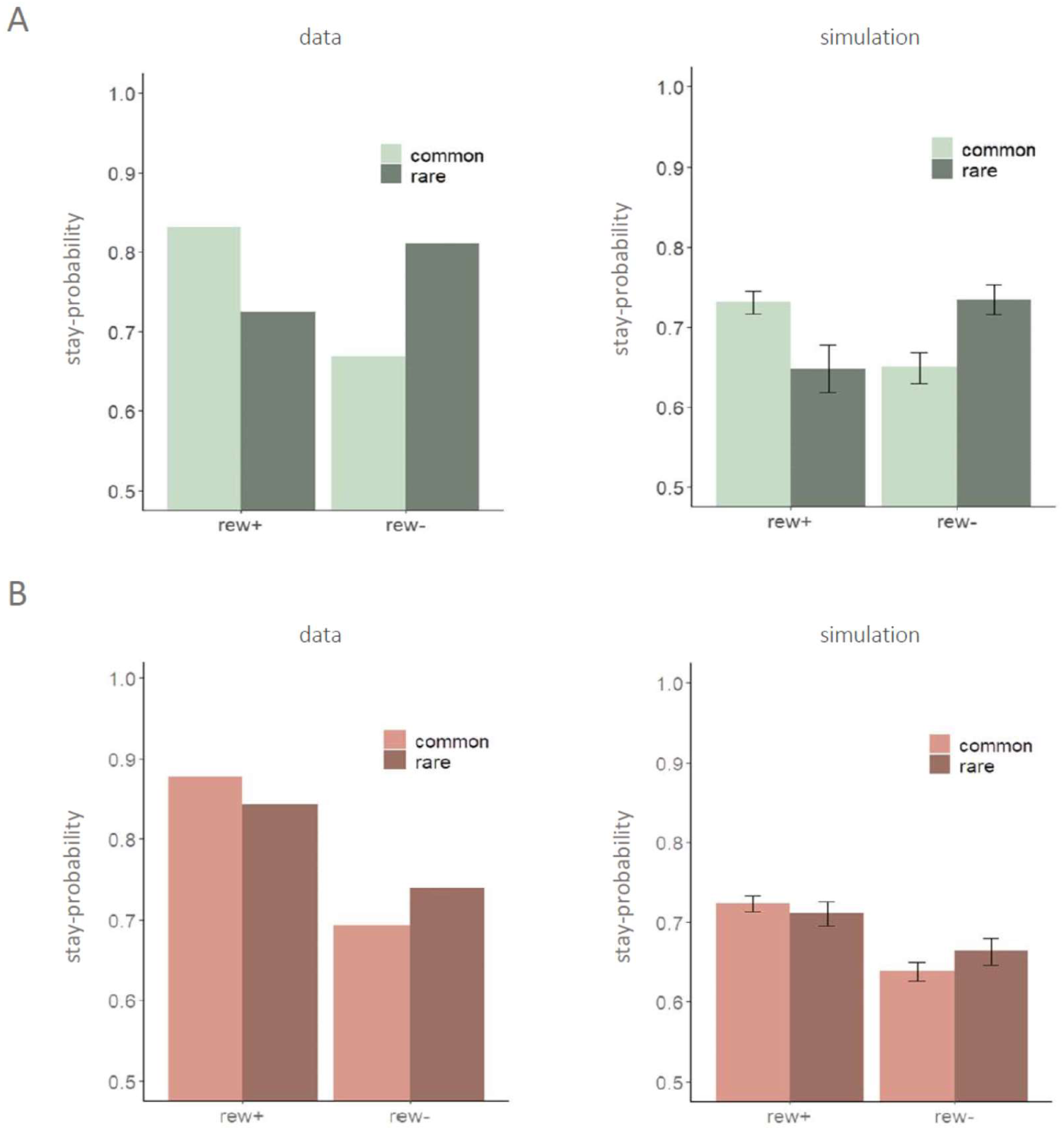
Probabilities of S1 choice repetition as a function of reward and transition type. Y-axis: Stay probabilities for 1st stage choices; data: empirical stay probabilities from data sets data1 (panel A; green) and data2 (panel B; orange). simulation: stay-probabilities from N=8000 simulated choice sequences per subject, derived from the winning model (M5).; rew+/-: previous trial was rewarded (+) or unrewarded (-).; common/rare: previous trial followed a common/rare transition respectively. Error bars in the simulation plots depict the 95% HDI over 8000 simulated data sets.

**Table 4.**
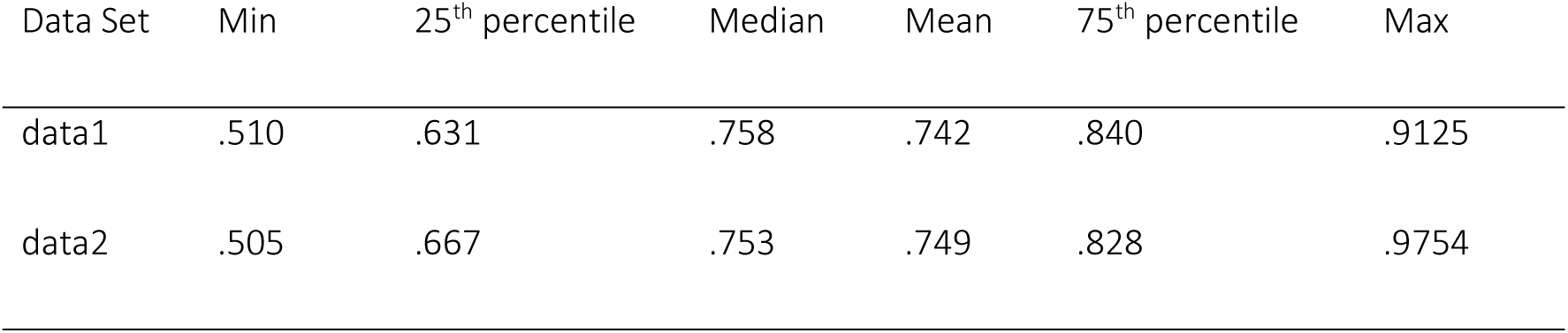
Proportion of correct S1 choice predictions by the winning model M5.

| Data Set | Min | 25<sup>th</sup> percentile | Median | Mean | 75<sup>th</sup> percentile | Max |
| --- | --- | --- | --- | --- | --- | --- |
| data1 | .510 | .631 | .758 | .742 | .840 | .9125 |
| data2 | .505 | .667 | .753 | .749 | .828 | .9754 |

#### Posterior Distributions

Figure 5 shows the posterior distributions of group-level parameters underlying S1 choices in the best-fitting model. The group-level mean of the exploration bonus parameter (ϕ) was positive in both data sets and the 95% highest-density interval (HDI) did not overlap with 0 (Table 5, Figure 5). We thus confirmed the predicted positive effect of directed exploration on S1 choice probabilities in both data sets. Furthermore, in both data sets, there was evidence for a substantial perseveration effect (Figure 5).

**Figure 5.**
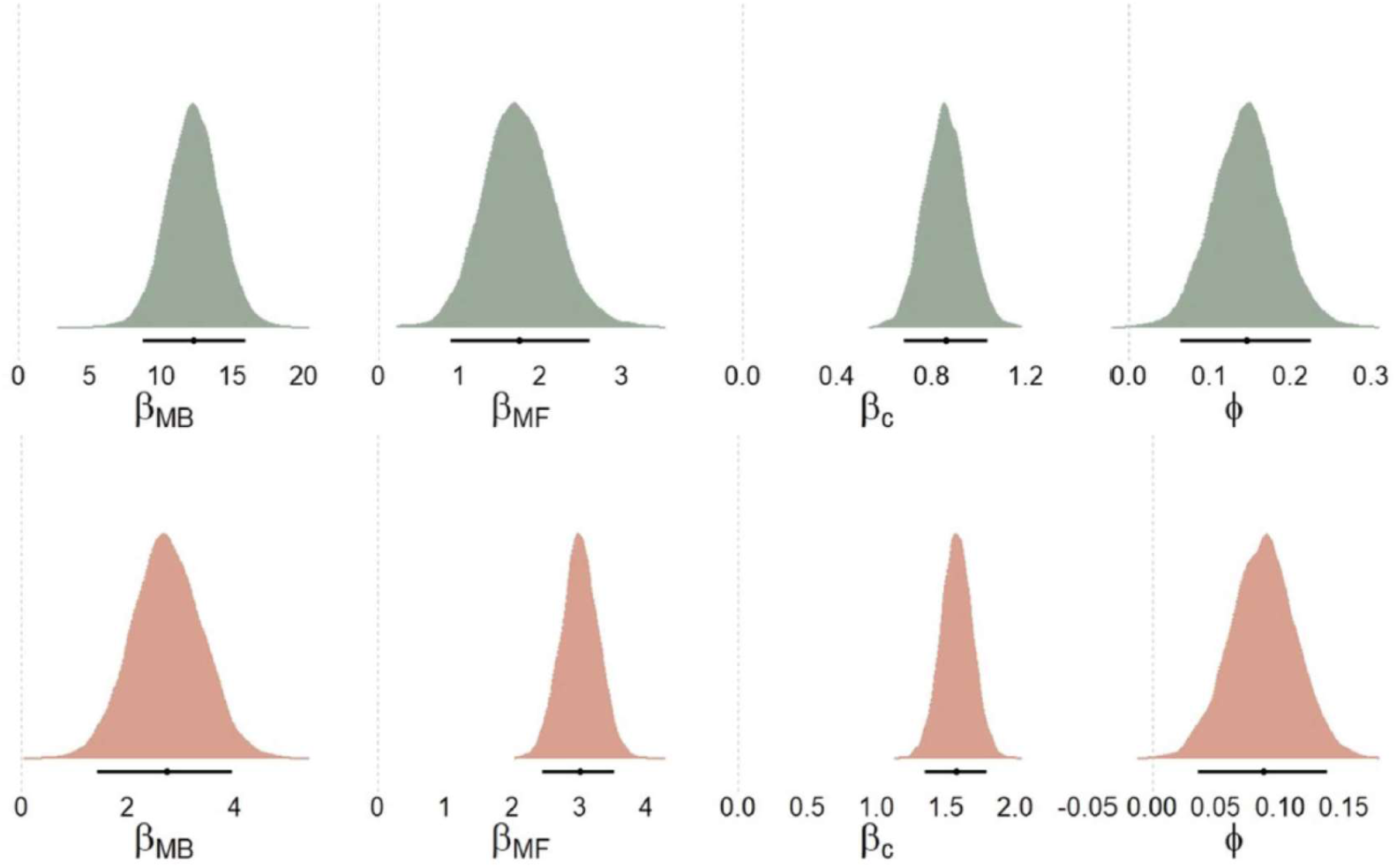
Posterior distributions of Group-Level Means of S1 Choice Parameters. Solid black lines show the 95% highest density interval (HDI) and the black dot depicts the point-estimate of the mean. Panels A and B (green and orange plots) show results on the basis of data sets data1 and data2, respectively.

**Table 5.**
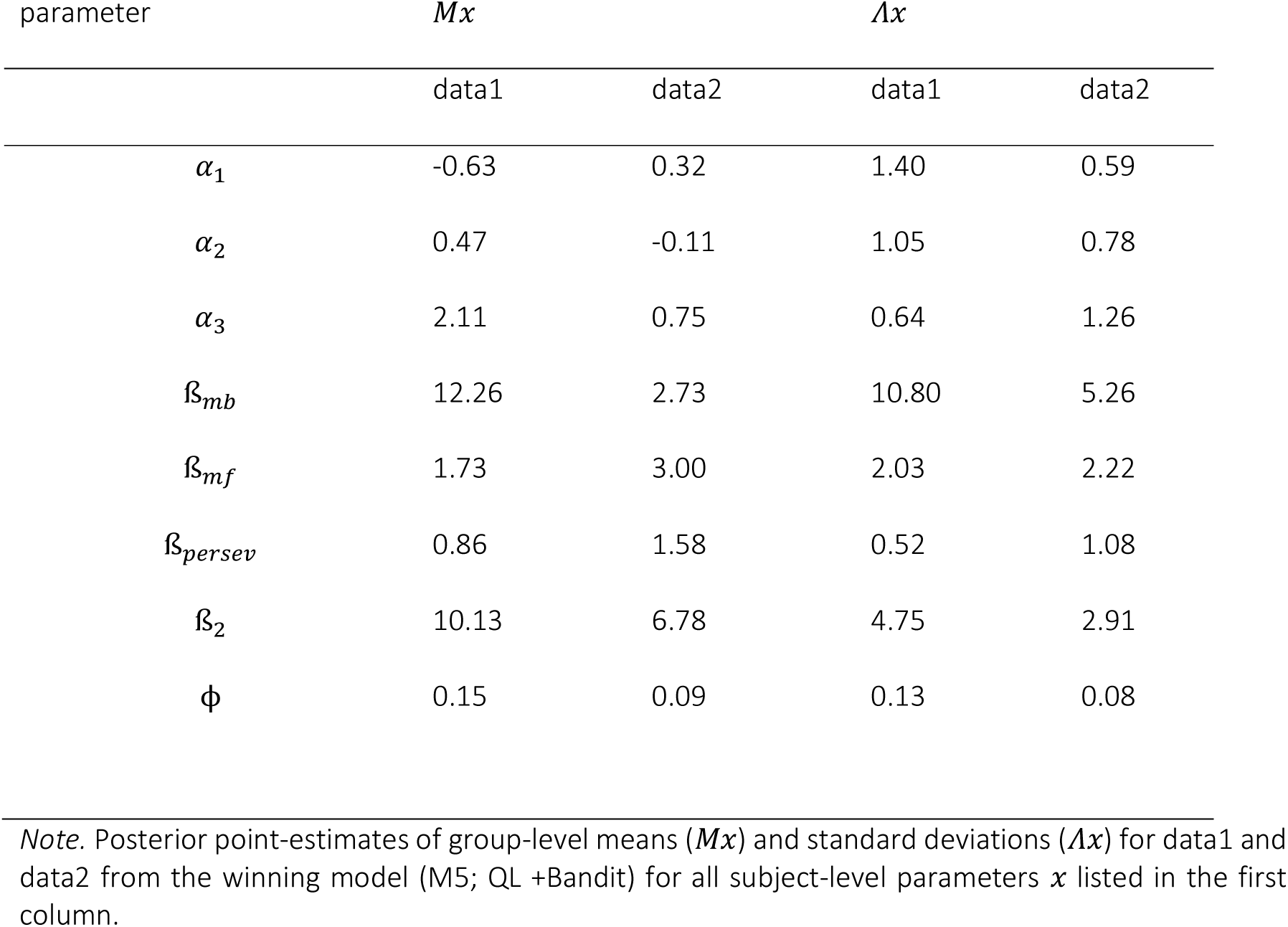
Posterior Point-Estimates of all Group-Level Parameters of model M5 for data1 and data2.

Posterior point-estimates of all hyperparameter means from the best-fitting model are shown in Table 5. Model parameters confirmed the model-agnostic analyses portraited above: the MB component (ß_mb_) was more pronounced in data1 vs. data2.

Recall that in the best-fitting model M5, the uncertainty heuristic (bandit heuristic) ranges between values of 0 to 3 per S2 option, resulting in a range of 1 to 5 for the combined predictor over both S2 options (see Equation 18 above). Recall further that rewards were re-coded to range between 0 and 1 for both data sets. Table 6 illustrates the resulting exploration bonus values (VEB) for four different example trials and parameter values of 𝜙 = 0.05; 0.1 and 0.15. Imagine a subject choosing option A in S1 (*S1A* first column), followed by a common transition, thus, leading to the S2-state A (*S2A* third column). Since the last visit of state S2A our subject may have chosen two and three alternative S2 options each, so that the *bandit-heuristic counters* of the currently available S2 options have values two and three respectively. Following our model, the sum of both counters (five in this example) is scaled by the transition probability (0.7 as it was a common transition from S1A) and the *alternative case* is added (see Equation 18). In this alternative case (following a rare transition, with probability 0.3 leading to S2B instead) *bandit counters* for the available choice options may have values zero and one respectively (e.g. if these were chosen in the two preceding trials). These resulting model-based uncertainty estimates are then weighted by the individual’s exploration parameter ϕ (here shown for exemplary values 0.05, 0.1, and 0.15).

**Table 6.** Exemplary exploration bonus values for different S1-S2 combinations given plausible exploration bonus parameter values for ϕ

| S1-choice | transition | S2 state<br>(*bandit* heuristic counter value) | $\phi$ | Resulting exploration<br>bonus value ($V_{EB}$) |
| --- | --- | --- | --- | --- |
| S1A | Common | S2A (2; 3) | 0.05 | 0.19 |
|  |  |  | 0.1 | 0.38 |
|  |  |  | 0.15 | 0.57 |
| S1B | Common | S2B (0; 1) | 0.05 | 0.11 |
|  |  |  | 0.1 | 0.22 |
|  |  |  | 0.15 | 0.33 |
| S1A | Rare | S2B (2; 3) | 0.05 | 0.115 |
|  |  |  | 0.1 | 0.23 |
|  |  |  | 0.15 | 0.345 |
| S1B | Rare | S2A (0; 1) | 0.05 | 0.17 |
|  |  |  | 0.1 | 0.34 |
|  |  |  | 0.15 | 0.6 |

*Note.* The first three columns show possible combinations of first state choices and second stage states following either transition type. $\phi$ : plausible exemplary subject-level parameter values. The last column shows the exploration bonus value for the these S1- S2 sequences for a given $\phi$ -estimate according to Equations 15 and 18.

A rough insight into the relationship of VEB compared to the weighted MB Q-value (QMB) of S1 options is shown in Table 7. In data1 the median proportion of VEB in relation to QMB (VEB / QMB over both S1 options) was 5%, whereas it was 17% in data2. These example calculations illustrate that, while posterior estimates of the exploration parameter may appear numerically small, they do in fact make meaningful contributions to subjective valuations.

**Table 7.** Proportion of VEB compared to QMB across both S1 options

| Data Set | 25<sup>th</sup> percentile | Median | 75<sup>th</sup> percentile |
| --- | --- | --- | --- |
| data1 | .03 | .05 | .11 |
| data2 | .08 | .17 | .42 |

*Note.* $V_{EB}$ and $Q_{MB}$ were calculated on the basis of subject-level parameter estimates derived from M5 (QL + Bandit). Values shown depict $V_{EB}/Q_{MB}$ (c.f. Eq. 18 and Eq. 7 respectively) over both S1 options, which were then averaged across all participants for data1 and data2 each.

#### Correspondence with model-agnostic analyses

In order to investigate how parameters derived from the model relate to model-agnostic indices of MB and MF behavior as well as to the overall performance, correlation analyses were performed (Figure 6). For this purpose, we applied the same regression model as described for the group analyses to each participant’s individual data set, omitting the random effects term, resulting in:

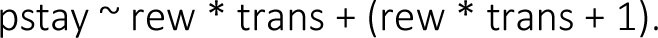

**Figure 6.**
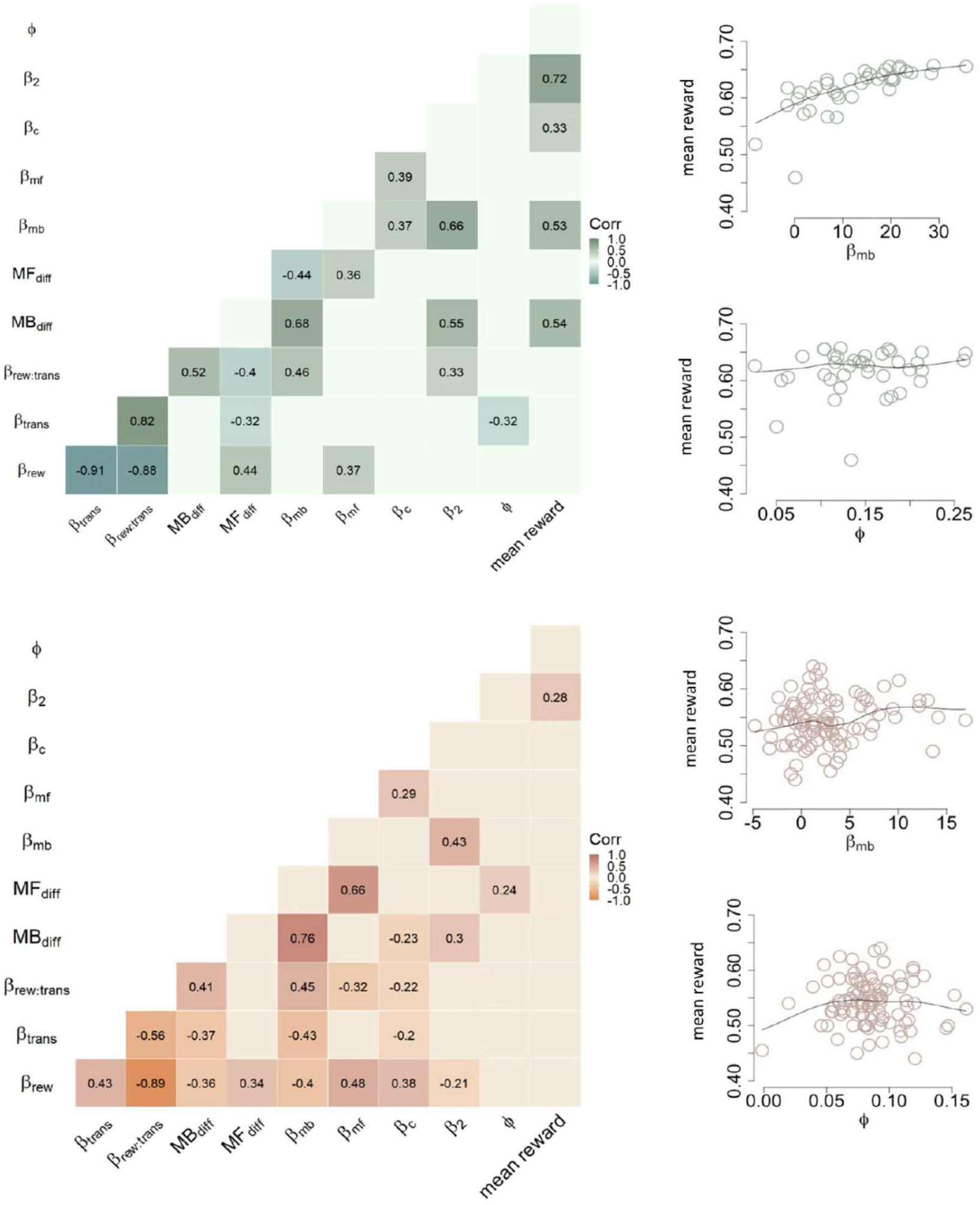
Associations of model-agnostic and model-derived TST indices of MB and MF signatures. Results from correlation analyses of model-agnostic indices of MB and MF influences. Empty tiles (left panel) indicate non-significant associations. ; ß_rew_,ß_trans_, ß_rew:trans_: regression weights for main effects of reward, transition type and their interaction; 𝑀𝐵_diff_, 𝑀𝐹_diff_: differences scores of MB and MF influences on S1 stay probabilities respectively; ß_MB_,ß_MF_: MB and MF S1 choice parameters from the winning model; ß_c_: S1 perseveration parameter; ß_2_: S2 inverse temperature parameter; 𝜙: exploration parameter; mean reward: mean reward gained throughout TST (data1: 300 trials, data2: 200 trials). Right panel: association of model-derived MB (ß_MB_) and exploration parameter (𝜙) with mean reward. Circles depict individual participants. All green plots (upper panel) are based on data1, orange plots (lower panel) are based on data2.

Model-agnostic indices of MB (ß_rew:trans_ and 𝑀𝐵_diff_) and MF (ß_rew_ and 𝑀𝐹_diff_) exhibited moderate associations in both data sets (Figure 6). Both effects were likewise associated with the corresponding model-derived parameters (ß_MB_, ß_MF_). The exploration parameter did not show significant associations with mean rewards earned on this task. In contrast, the MB component showed a moderate to strong association to the mean overall payout in data1 but not data2, confirming that the modified version successfully addressed previous concerns (see Kool et al., 2016). In data2, there was no evidence for this association (see Figure 6, lower panel).

### Further Validation of the best-fitting model

In a final step, we verified that results for data2 were not due to specific characteristics inherent to the subsample we drew from the data set from Gillan et al. (2016). To this end, we repeated the model estimation procedure for M5 (c.f. Methods described above) using the full sample of experiment 1 (N=548) from the original publication by Gillan and colleagues (2016). Modelling results were comparable to those reported above for the initial subsample (data2). The model converged equally well (all 𝑅^-^<1.1) and posterior estimates of group-level parameters mirrored results reported above (Figure7). Results again confirmed a highly robust effect of S2 uncertainty on S1 choice probabilities (see Figure 7), such that the posterior distribution of the exploration bonus parameter 𝜙 was robustly > 0.

**Figure 7.**
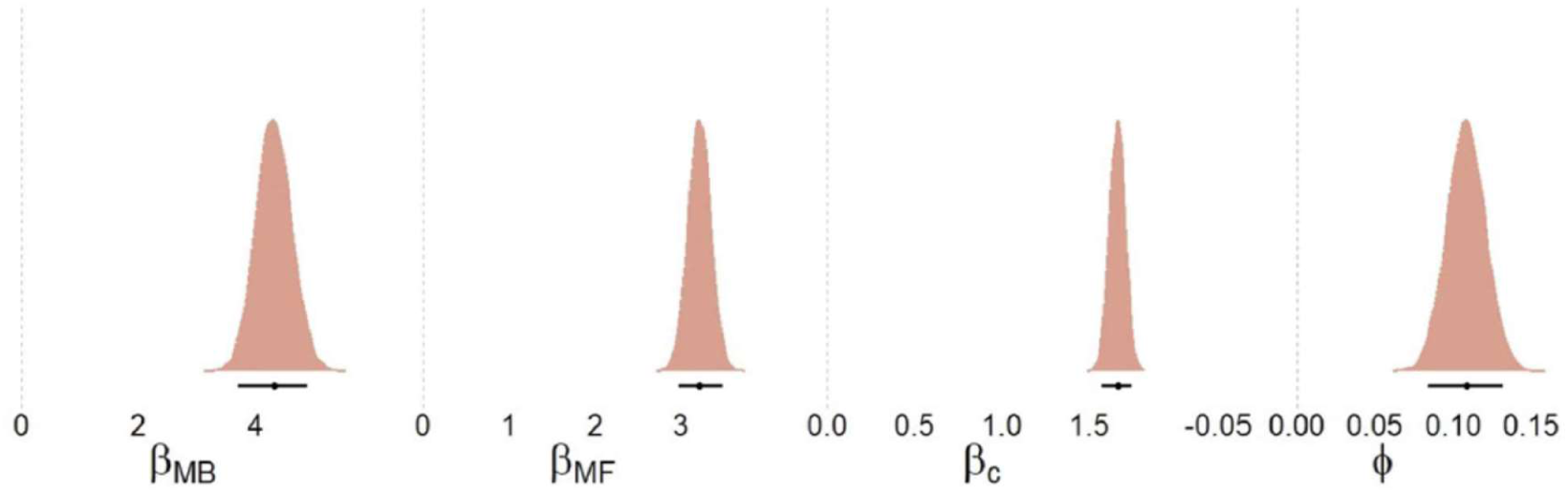
Posterior density estimates of 1^st^-stage group-level mean choice parameters from the winning model. The lower panel of Figure 5 shows corresponding results based on data2 (N=100), plots shown here are based on the full sample of experiment 1 in the original publication (N=548; Gillan et al., 2016). Black dots indicate the mean point-estimate, black bars cover the 95%-HDI.

## DISCUSSION

This study set out to extend existing hybrid models of TST behaviour with mechanisms implementing directed (uncertainty-based) exploration. To this end, we considered varying ways in which uncertainty may guide participants choices in stage 1 of the task, building upon insights from the explore-exploit dilemma (4-armed restless bandit; Daw et al., 2006; Chakroun et al.,2020; Wiehler et al., 2021). In the empirical data analysed here, an uncertainty-dependent learning model (BL-models using a Kalman Filter) did not provide an advantage over a classic Q-Learner algorithm. The addition of heuristic-based uncertainty measures in the choice component for stage 1 of the task, however, significantly improved model fit (see model comparison results). Results were consistent across two independent data sets using two versions of the TST assessed in different settings (laboratory vs. online sample), and confirmed a positive directed exploration effect in TST behavioural data.

We complemented model-based results with more traditional model-agnostic analyses to gain a deeper understanding of how these relate to each other. This additionally aided the interpretation of the best-fitting model, parameter estimates derived from it as well as differences in the results obtained from both data sets.

### Uncertainty-based exploration (but not learning) on the TST

While human choice behaviour in the bandit task is typically better described by models that incorporate a Kalman-Filter as a learning component (vs. a constant learning rate; see e.g. Daw et al., 2006; Raja Beharelle et al., 2015; Chakroun et al., 2020; Wiehler et al., 2021), this was not the case for TST data analysed here. This may likely be due to the more complex task structure and resulting higher cognitive demands. Thus, tracking the underlying reward walks and uncertainties associated with them might be too computationally demanding. Participants did, however, show uncertainty-dependent adjustments in their choices in S1 of the task, such that robust exploration bonus parameters > 0 were obtained (see below). Yet, when faced with a more complex task, participants might fall back on simpler, heuristic-based uncertainty proxies to guide their behaviour. In fact, the current results point to an advantage of the more parsimonious counter-based heuristic tested here (M5, see model comparison results in favour of the bandit-heuristic compared to the trial-heuristic model M7). Although a purely temporal indicator (M6 & M7, trial heuristic) may seem more intuitive at first, continuous tracking over the course of up to 300 trials (data1; 200 trials in data2) seems cognitively implausible. The bandit heuristic further has the advantage of visual support throughout the task (clearly distinguishable stimulus identities). These external cues, alongside the limited numerical range, ease the demand placed on working memory processes and thus may ultimately “protect” the MB system (Otto et al., 2013; Brown et al., 2022; Collins et al., 2017; Yoo & Collins, 2022; for a diverging view however see Silva et al, 2022). As the instructed goal for participants was to maximize their pay-outs, placing higher priority on the MB system might be advantageous. As Kool and colleagues (2016) have pointed out, such an advantage is however, dependent upon the task version at hand. This finding is supported by the significant positive association of MB control and rewards earned in data1 (adapted version of the TST) and a lack thereof in data2 (classic TST version). These differential associations also show that previously voiced criticisms and proposed alterations (Kool et al., 2016) have successfully been addressed in the adapted task version used in data1. Parameter estimates for the weight on 2^nd^-stage Q-values were positively associated with overall pay-out in both data sets and were indeed the only significant association in this regard in data2. Thus, relatively lower MB tendencies in data2 may even be seen as goal-directed in a broader sense, as MB control is commonly viewed as more demanding, while in this case not more rewarding and ultimately too costly.

Taking a closer look at the exploration bonus from the winning model, we show that here too, results from data1 and data2 differ in regard to their association with MB control. Table 7 provides rough estimates of the relative proportion of this bonus in relation to MB control. As pointed out above, MB control was attenuated in data2 compared to data1. Considering the similar posterior estimates for the exploration parameter these proportional differences (VEB / QMB : 5% vs 17% of in data1 and data2, respectively, Table 7) come to little surprise. Nonetheless, one may speculate that they are not fully independent of aforementioned diverging utility and contribution of MB control. Varying reward magnitudes in the adapted TST version (data1) are likely easier to track than binary outcomes in data2 and may have provided participants with a sufficiently useful and not overly costly reference point for S1 preferences. Tracking underlying reward dynamics based on binary outcomes (data2) on the other hand might be more difficult. The additional heuristic-based uncertainty estimates may have thus been of varying utility during decision-making, depending on the environment (i.e. task version) at hand. These considerations are somewhat speculative in nature and warrant further, more detailed investigations in the future.

The assessment within a laboratory setting further enabled more control over participants’ understanding of the task at hand compared to the online sample from which we derived data2. As several scholars have pointed out over the past years, general task understanding and diverging instructions of the TST can have a significant influence on the relative employment of the MB over MF system (e.g. Akam et al., 2015; Feher da Silva & Hare, 2018; Castro-Rodrigues et al., 2022; Hamroun, Lebreton,& Palminteri, 2022). It should be noted that the authors of the original publication of data2 have taken extensive precautions such as training trials and a comprehension test (for details see Gillan et al., 2016) prior to execution of the TST. Nonetheless, insight into participants’ model of the task remains reduced within this online context. Noticeable differences in the compensation and therefore incentive for participation may have further influenced individuals’ motivational state with regard to more effort placed on task execution (see e.g. Patzelt et al., 2019).

### Dual-System Views and the TST

Beside these external influences on relative MB and MF contributions (i.e. task versions, instructions, incentives etc.) broader criticisms regarding their definition within classic dual-system frameworks has been raised. As outlined previously, indices of MB control likely only depict one possible goal-directed strategy subjects use to complete the TST (recall reduced MB control in data2 vs. data1 in light of its utility for reward maximization, i.e. long-term goal-attainment). Consequently, several alternative strategies that may also utilize a model of the environment, rendering them MB in the literal sense, are not accounted for (Feher da Silva & Hare, 2018; 2020; Toyama et al., 2017;2019). Models employed on the other hand, may be skewed, outright incorrect, or employed in a rigid and habitual way, further complicating a clear-cut interpretation of associated indices (see e.g. Seow et al., 2021; Shahar et al., 2019a). The same holds true for potential additional subprocesses which are not represented in classic dual-system views and formalizations thereof (Collins et al., 2017; Collins & Cockburn, 2020; Feher da Silva et al., 2022). At this point it should be noted that several of these issues may also apply to the explore-exploit research and theoretical assumptions works in this field are based on.

To address these concerns several scholars have been developing adapted versions or novel alternatives to these paradigms (see e.g. Kool et al., 2016 and the adapted TST version applied by Mathar et al., 2022; Bruder et al., 2021; Wagner et al., 2021 etc.). To name one prominent example posed as an alternative (or at the least useful supplement) to widely applied classic restless bandit paradigms in the explore-exploit research, Wilson and colleagues (2014) have introduced the *Horizon Task.* This task aimed at the decoupling of reward and information, which are classically confounded and thus hamper the clear distinction between exploration, exploitation and their driving factors. The Horizon Task has been successfully applied in a number of studies and has thus far also undergone several further adaptations (e.g. Feng et al., 2021; Cogliati Dezza et al., 2017; Sadeghiyeh, et al., 2020).

### Computational Modelling

Another approach is the development of novel computational models to better delineate the specific processes engaged during task performance. Several researchers have set out to move away from traditional computational accounts, and build upon markedly different conceptualisations of learning and decision processes in this task. One recent example for such efforts comes from Gijsen, Grundei, and Blankenburg (2022): The authors applied an active inference account to TST data and – akin to the procedure laid out here-re-analysed existing data sets. Some of these were in fact better described by the proposed more elaborate models. However, specifically data gathered in an online setting as well as data including a negative reinforcement scheme diverged with regard to model ranking. The preferred model for these data sets (referred to as *online* and *shock* data sets respectively in Gijsen et al., 2022) was in fact more akin to versions tested here. Moreover, it should be noted that the current study followed a different aim in more general terms. As pointed out previously, the TST is widely applied and probably the most utilized paradigm to capture MB and MF behavioural tendencies.

Thus, by extending existing models that are already in use, we hope to balance improving their descriptive ability while at the same time ensuring their applicability for a wide scientific audience. By making the code for the winning model freely available we hope to foster similar efforts (c.f. *Open Code* above).

### Limitations of the current study

Common to all computational modelling approaches are basic considerations regarding the limited scope of possible mechanisms accounted for. To put it more bluntly: Results derived from computational models (and their comparison) are ultimately limited to the finite set of processes defined in them. Due to its non-specific applicability this issue may almost seem trivial, but should nonetheless be kept in mind when evaluating and interpreting such results.

In addition to these broader conceptual issues with regard to the TST, more specific limitations of the present study should be noted as well. While results from model comparison clearly favoured all QL over BL models in both data sets it should be noted that the implemented belief updating process in the latter model group depicts an approximation (vs. exact representation) of the true underlying random walk dynamics (control analyses however showed that the empirical reward dynamics closely corresponded to those implemented in all BL models).The Kalman-Filter updating process between trials (Equation 12) in all BL models is analogous to the *forgetting* process implemented in the QL model variants (Equation 4). Thus, for both learning mechanisms we assumed subjective value estimates of unchosen options to move closer to a reasonable estimate (mid-range of possible values). Nonetheless, future applications may consider refining this model aspect.

Despite the fact that the evidence for strategic exploration behaviour was consistently present across both data sets (exploration bonus effects were substantially larger than zero), these effects were numerically small (e.g. mean exploration bonus values ranged between 5% and 17% of MB values). Yet, inclusion of these terms substantially improved model fit. As outlined above, model variants shown in the present study only cover a very limited set of possible exploration mechanisms at play. Different versions of such exploratory behaviour or additional processes not accounted for here are possible, in light of these results plausible or even likely and should be part of future investigations.

These may also include simulations and analyses thereof in order to provide comparison results to those presented here. To our knowledge such analogous simulations from related work are currently not available. Thus, a clear interpretation of the winning model’s simulation presented above is hard to reach. Although posterior predictive checks revealed that the winning model (M5) accounted for the over data pattern quite well, it still underpredicted S1 stay-probabilities in both data sets (c.f. Table 4 and Figure 4 above). Future work is required to determine the degree to which this depends on the specific task version employed, or reflects a general shortcoming of current hybrid models.

Another potential limitation is that recently applied DDM choice rules were not used in the present study (Pedersen, Frank, & Biele, 2017). The investigation of reaction time distributions and their relation to information processing and decision-making can provide valuable insights that may complement present results (Shahar et al., 2019b). Parameters derived from models like these have further been linked to various (sub-)clinical symptoms, and thereby also shed light on potential disease mechanisms (see e.g. Forstmann, Ratcliff, & Wagenmakers, 2016; Mandali et al., 2019; Maia, Huys, & Frank, 2017, Sripada & Weigard, 2021). Here however, our goal was first to confirm the existence of the proposed exploration mechanisms in different TST versions. We therefore leave extensions to DDM choice rules to future work.

The empirical data used to develop and test the proposed novel model variants may pose yet another concern. We included two TST versions as a first step, but several additional task variants could be examined in future work in order to further validate the adapted hybrid model. Considering aforementioned ambiguities with regard to the generalizability and transferability of proposed exploration-strategies (and other information-processing steps inherent in the model; e.g. learning mechanisms etc.) these may also be tested for in data from related paradigms posing similar demands (i.e. reward accumulation in light of uncertain, dynamic environments). In order to (at least in part) account for the myriad of contextual factors influencing learning and decision-making in such paradigms, data from within-subject design studies examining specific contextual effects would be required to delineate how the proposed exploration mechanism is modulated by these factors.

An additional issue concerns the generalizability of the present results. To this day, a large part of empirical findings is based on small rather homogenous groups of individuals, namely WEIRD ones (i.e. white, educated, industrialised, rich, & democratic), which also applies to many other data sets in the field. Despite participants’ *WEIRDness*, samples are seldomly diverse with regard to age or gender either. In the present case for example, the sample from data1 was exclusively comprised of 18 to 35-year-old heterosexual males (Mathar et al., 2022). While reducing variability in these sample characteristics has its’ utility, (improving internal validity and thus enabling more clear-cut interpretations) results are consequently limited to this confined group. Gillan and colleagues (2016) on the other hand employed a more diverse large-scale community sample. Despite lesser concerns regarding diversity, here other limitations that arise due to the online setting and associated factors come into play (e.g. data quality due to false profiles, low incentives, task understanding etc.).

Despite ever growing popularity and application of transdiagnostic as well as dimensional conceptualisations of mental health, a substantial body of research is still based on the comparison of groups defined as either *healthy* or *diseased.* Again, procedures like this have a rational basis, entail advantages, and have produced a wealth of valuable insights. Keeping this and aforementioned progress in mind (Insel et al., 2010; Robbins et al., 2012; Maia & Frank, 2011), it is nonetheless warranted to push further. Future studies that leverage large samples and the natural sub-clinical variation in psychiatric symptomatology these entail, are called for.

## CONCLUSION

In summary, we provide computational evidence for a contribution of directed exploration to behaviour in two independent data sets of different variants of the a widely used two-step task (TST). We compared a series of extensions of commonly applied hybrid models for TST behaviour using concepts from exploration-exploitation work (Wilson et al., 2021), and show that a model with a heuristic-based directed exploration term for S1 decisions consistently outperformed both a baseline model without a directed exploration process, and a series of alternative model variants. Future work may extend these approaches to other task variants, and explore the degree to which the observed directed exploration process observed for TST behaviour is sensitive to e.g. individual differences in (sub-)clinical measures of psychopathology.

## Author contributions

A.M.B. and J.P. developed the model extensions. D.M. provided analytical tools. A.M.B. analysed the data and wrote the paper. J.P. and D.M. provided comments and revisions.

## Acknowledgements

This work was supported by a grant from Deutsche Forschungsgemeinschaft (DFG) (PE1627/5-1 to J.P.)

## SUPPLEMENT

**Figure S1.**
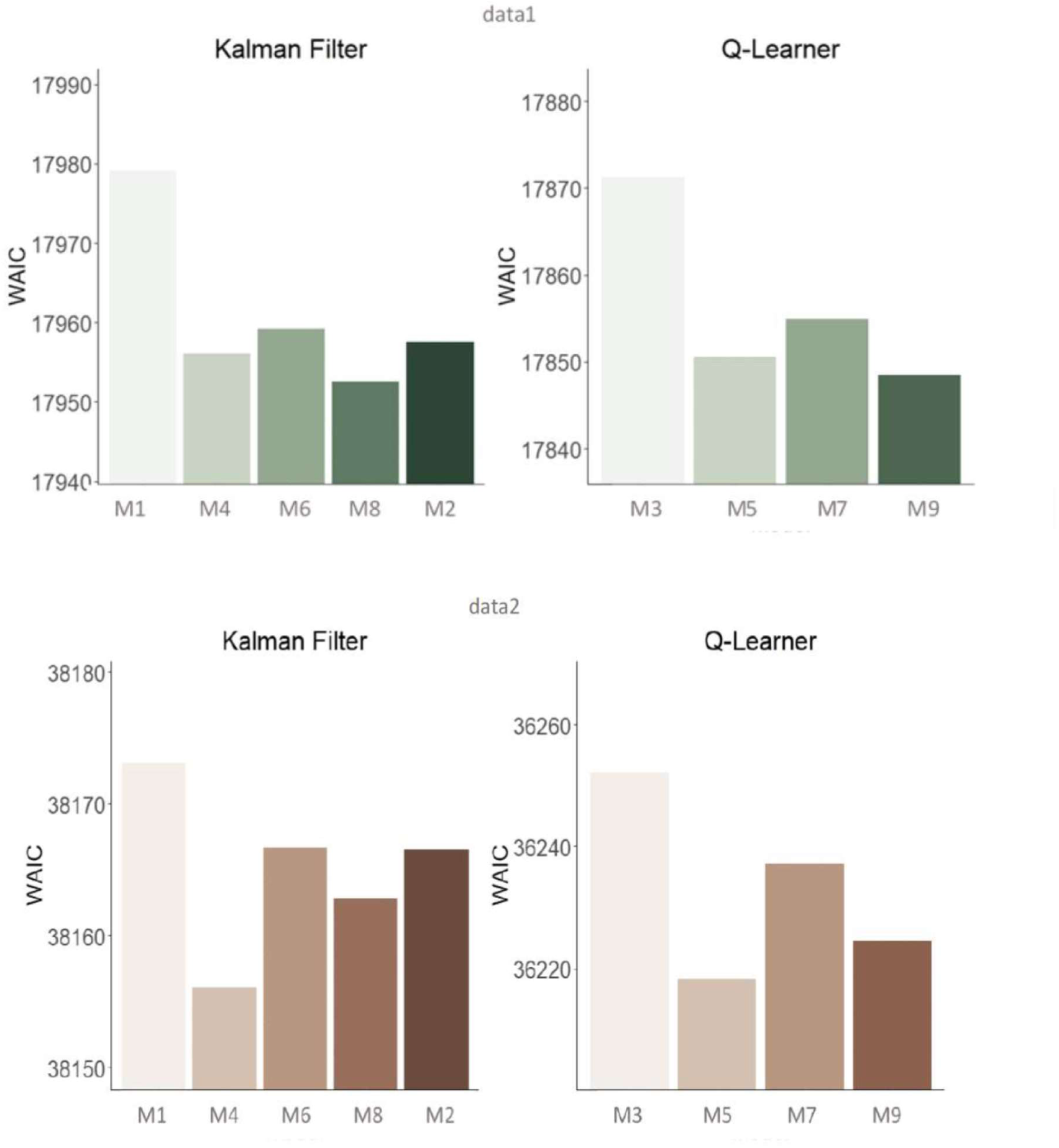
Model Comparison Results for all models considered via the Widely Applied Information Criterion (WAIC). Panel A/B green/orange bar plots refer to data1 and data2, respectively. Kalman-Filter: Bayesian Learner (BL) models.

